# Long-term single-cell passaging of human iPSC fully supports pluripotency and high-efficient trilineage differentiation capacity

**DOI:** 10.1101/663047

**Authors:** Estela Cruvinel, Isabella Ogusuku, Rosanna Cerioni, Jéssica Gonçalves, Maria Elisa Góes, Juliana Morais Alvim, Anderson Carlos Silva, Alexandre Pereira, Rafael Dariolli, Marcos Valadares, Diogo Biagi

**Author notes:** Both authors contributed equally to this work (Estela Cruvinel, Isabella Ogusuku). Mailing address: 2242 Prof. Lineu Prestes Avenue, Sao Paulo, SP. 05508-000, Brazil.

## Abstract

**Objectives:** To validate a straightforward single-cell passaging cultivation method that enables high-quality maintenance of hiPSC without the appearance of karyotypic abnormalities or loss of pluripotency.

**Methods:** Cells were kept in culture for over 50 passages, following a structured chronogram of passage and monitoring cell growth by population doubling time (PDT) calculation and cell confluence. Standard procedures for iPSC monitoring as embryonic body (EB) formation, karyotyping and pluripotency markers expression were evaluated in order to monitor the cellular state in the long-term culture. Cells that underwent these tests were then subjected to differentiation into keratinocytes and cardiomyocytes to evaluate its differentiation capacity.

**Results:** hiPSC clones maintained its pluripotent capability as well as chromosomal integrity and were able to generate derivatives from the three germ layers at high passages by embryoid body formation and high-efficient direct differentiation into keratinocytes and cardiomyocytes.

**Conclusion:** Our findings support the routine of hiPSC single-cell passaging as a reliable procedure even after long-term cultivation, providing healthy PSCs to be used in drugs discovery, toxicity and disease modeling as well as for therapeutic approaches.

## INTRODUCTION

Human Induced Pluripotent Stem Cells (iPSC) unique features make them reason for huge excitement among scientific and medical communities as an alternative to human embryonic stem cells (hESC) for research on tissue-specific development, disease modeling, cell-based therapies and drugs discovery. Since iPSC were reported,^1^ culture methods have evolved to optimize growth conditions while maintaining pluripotency in order to meet both the special needs of the cells and their rapid demand. However, many laboratories still cultivate iPSC following out-of-date procedures that remain from the discovery of hESC such as colony passaging,^2^ which is an approach that usually follows uneven confluency as well as unpredictable growth rates.^2,3^

To overcome colony passaging-related issues, several groups committed to the search of conditions that could support single-cell passaging,^4–8^ but despite the rapid progress there is still a lack of standard protocols for iPSC cultivation that comprises the effects of combining different biotechnologies as well as the effects when culturing the cells for a long time. Several recent reports present lack of information concerning iPSC genetic integrity,^9^ especially after long-term cultivation *in vitro*,^10^ even though many studies suggest abnormalities to be progressively favored by suboptimal culture conditions such as single-cell passaging^11^ or high-cell density cultures.^12^

It is common for laboratories to maintain multiple iPSC cell lineages in culture to avoid the inconsistencies of long-term cultivation, however it is important to recognize that this approach implies high cost and labor demand and also does not guarantee success for differentiation protocols, as working with different clones impairs reproducibility, homogeneity and scalability in differentiations.^13^ Nevertheless, as long-term maintenance in culture seems likely to promote self-renewal^10,14^ and limit differentiation through progressive selection of genetic variants,^15^ the assessment and validation of iPSC cultivation protocols is mandatory to support reproducibility in differentiations.

In the present work we used hiPSC derived from two distinct primary sources to develop a controlled long-term culture methodology that supports single-cell passaging while maintaining pluripotency markers, genomic integrity and high ability to generate derivatives by directed differentiation, resulting in high-purity specialized cells.

## METHODS

### Ethics Statement

This investigation agrees with the principles outlined in the Declaration of Helsinki and the study protocol was approved by the Ethics Committee for Medical Research on Human Beings of the Institute of Biomedical Sciences from the University of São Paulo (#2.009.562). Signed informed consent was obtained from all participants.

### iPSC Reprograming and Maintenance

Human erythroblasts and skin fibroblast were reprogrammed for human induced pluripotent stem cells generation (hiPSCs). Erythroblasts were genetically modified using an episomal vector system previously described in the literature ^16,17^ with slight modifications. Briefly, mononuclear cells were isolated from 10□mL of peripheral blood by Ficoll gradient. Erythroblasts were cultured in a serum-free mononuclear cell (MNC) medium containing the following cytokines diluted in Stem Span Serum Free Expansion Medium (Stem Cell Technologies, CA): 40□ng/mL insulin-like growth factor 1 (IGF1); 100□ng/mL Stem Cell Factor (SCF); 10□ng/mL interleukin 3 (IL-3); 2□U/mL erythropoietin (EPO). Cells were transfected with plasmids pEB-C5 and pEB-Tg (Addgene, USA) containing reprogramming factors OCT4, SOX2, KLF4, cMYC, LIN28 and SV40-T, using the Human CD34+ nucleofector kit and the Nucleofector II device (both from Lonza Group Ltd., CH) following manufacturer’s instructions. Reprogrammed erythroblasts were incubated in MEF-coated plates in MEF medium and FBS ES-Cell Qualified (ESQ; Thermo Fisher, USA) with 20□ng/mL basic fibroblast growth factor (bFGF) overnight. Then, they were transferred into embryonic stem cell (ESC) medium containing Knockout DMEM, Knockout Serum replacement, antibiotic-antimycotic, 200□mM Glutamax, MEM non□essential amino acid solution and 2□mercaptoethanol (all from Thermo Fisher, USA) supplemented with 20□ng/mL bFGF and 0.25□mM Sodium butyrate (Sigma Aldrich, EUA). hiPSC colonies were collected from a 6 well MEF-coated plate using Gentle Cell Dissociation Reagent (Stemcell Technologies, CA) and seeded into Matrigel (BD, USA) coated plates with Essential 8™ medium (E8; Thermo Fisher, USA) supplemented with 10◻μM ROCK inhibitor Y-27632 (Stemgent Inc., EUA).

hiPSC lines were derived from fibroblast following Epi5™ Episomal iPSC Reprogramming Kit protocol (Invitrogen) with some modifications. Briefly, fibroblasts were isolated from skin biopsy of a healthy donor by manually processing into smaller pieces using a scalpel. Fibroblasts were cultivated in DMEM High Glucose medium supplemented with 15% bovine fetal serum, 200□mM Glutamax, 100 U/mL penicillin and 100 μg/mL streptomycin (all from Thermo Fisher, USA). Fibroblast were reprogrammed at 3rd passage and 2.7×10^4^ cells were platted in GELTREX™ matrix coated wells from a 12-well plate. In the next day, Epi5™ Episomal iPSC Reprogramming Kit factors were added in the cell using OptiMEM medium and Lipofectamine 3000 reagents (all from Thermo Fisher, USA). Reprogrammed cells were maintained with Essential 6™ medium (Thermo Fisher, USA) supplemented with 100 ng/mL bFGF (R&D Systems, EUA) and 100 μM of sodium butyrate (Sigma Aldrich, EUA) for 14 days, then the medium was switched by E8 medium and 100 μM of Sodium butyrate for more 14 days. From the 14th to 28th day after reprogramming small colonies emerged and were manually picked and seeded individually into new GELTREX™ matrix coated wells. After the 28th day, cells were maintained with E8 medium.

Blood and skin samples were collected from four healthy donors ranging in age from 30 to 40 years old: two males (ACP, PC3) and two females (PC2, PC4). From these samples four cell lines were generated being one derived from erythroblasts (ACP) and three derived from skin fibroblasts (PC2, PC3, PC4). After clonal picking and expansion, one clone of ACP (ACP5) and three clones of the lines PC2 (PC2.2, PC2.3, PC2.4), PC3 (PC3.1, PC3.2, PC3.3) and PC4 (PC4.3, PC4.5, PC4.6) were selected. Due to their growth rates and the fact they were obtained by using different sets of plasmids for reprogramming thus allowing more interesting comparisons, only ACP5 and PC4 clones (PC4.3, PC4.5, PC4.6) were used for the next steps of long-term cultivation, characterization and differentiation.

From passage 5 and forward, we followed the method described in Figure 1–A. Briefly, iPSCs were passaged twice a week after three and four days of culture. Cell density determination will be explained bellow. iPSCs were cultivated with E8 medium and Essential 8 Flex medium™ (E8 Flex) over a GELTREX™ matrix (all from Thermo Fisher, USA). Passages were performed by incubating the cells with Versene™ supplemented with 10% Triple Express (both from Themo Fisher, USA) for 5-7 minutes and gentle pipetting for colony dissociation, cells were then centrifuged 4 minutes at 300xg and resuspended in E8 medium with 10 nM Y-27632 (Cayman, USA). hESC BR1 (donated by Prof. Dr. Lygia V. Pereira^18^) was also cultivated as described above and used for comparative analyses when appropriate.

**Figure 1.**
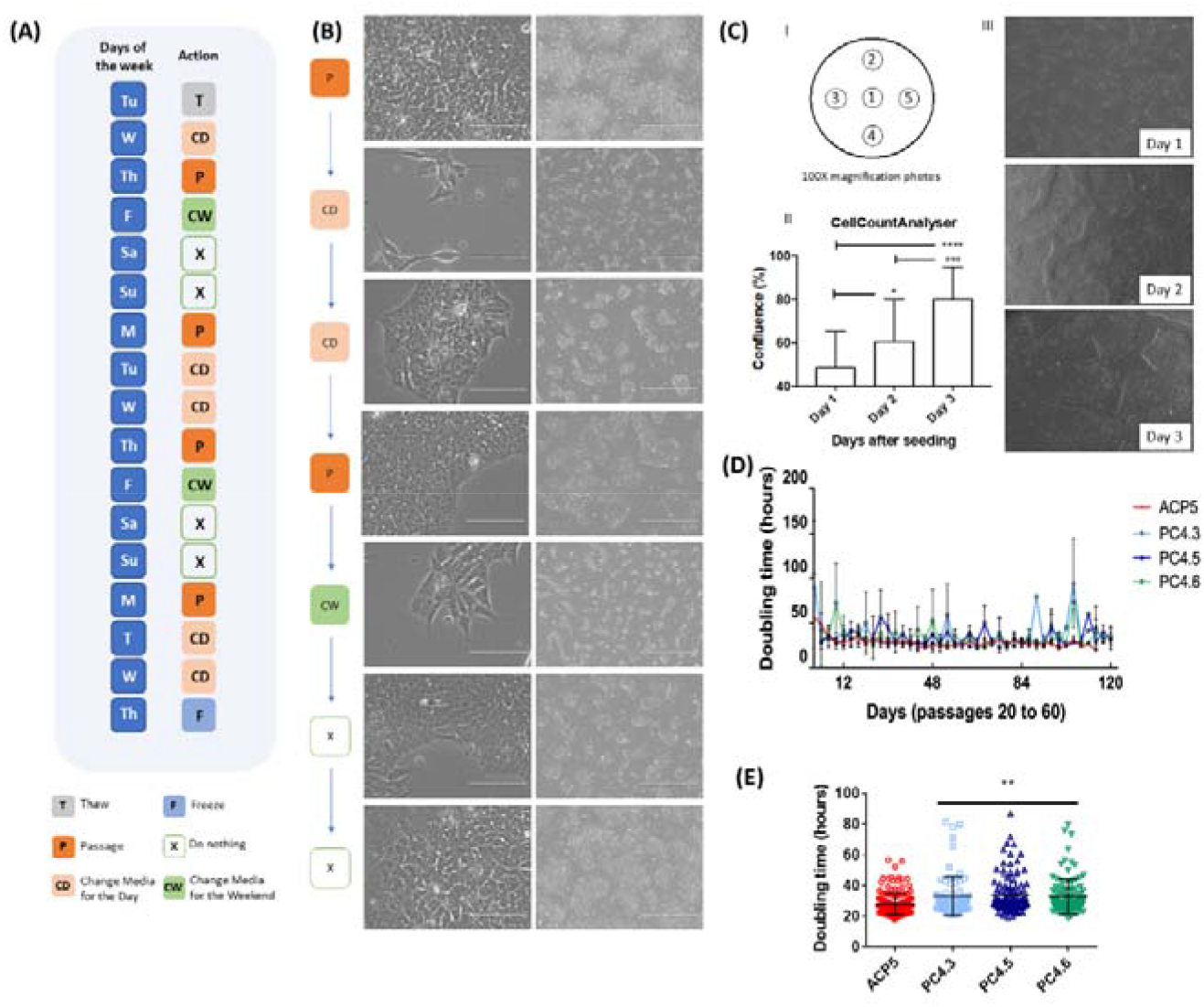
Single-cell passaging did not impair hiPSC morphology or colony formation capacity. **(A)** Scheme of the hiPSC maintenance workflow for the week. **(B)** Cells morphology throughout the days before media changing. Right line scale bar represents 1000 μM and left line scale bar represents 100 μM. **(C)** Cell confluence was monitored by CellCounterAnalyser. I) Representative scheme of the 5 recorded photos per 6 wells plates, II) quantification of 5 different passages recorded 3 days after seeding, data presented as mean ± SD. *p<0.05; ***p<0.001 and ****p<0.0001 by ANOVA, III) representative images of the photos recorded for quantification. **(D)** PDT of hiPSC clones ACP5 (n=8), PC4.3 (n=5), PC4.5 (n=4) and PC4.6 (n=5) from passage 20 to 60**. (E)** All PDT values obtained for clones ACP5 (n=165), PC4.3 (n=83), PC4.5 (n=100), PC4.6 (n=112). Data is presented as mean ± SD. **p< 0.01 vs. ACP5 by ANOVA.

### Population doubling time calculation

For population doubling time (PDT) calculation, hiPSC were counted first when seeding the cells and then when detaching them for passage. At each time point cells were counted twice with Trypan blue using a Neubauer’s chamber and mean was used to determine total number of cells. PDT calculation was determined by PDT= t. Log10(2)/ [Log10(x) – Log10(x0)], whereby t is expressed in hours. Results were then plotted as mean ± standard deviation (SD) for each given day. The significance of differences among passages was analyzed for each clone through ANOVA and P <0.05 was considered statistically significant.

### Confluence monitoring – CellCountAnalyser software

We used an ImageJ associated software previously published by Busschots et. al.^19^ and upgraded for our team (and so renamed CellCountAnalyser; details can be found in the supplementary material) to monitor and work only with cells in logarithmic phase of growing. To do so, 24-96 hours after cell seeding, five photos were recorded from marked areas that covered center and borders of cell dishes using EVOS FL Optical microscope (Thermo Fisher, USA). Then using our one-click Python-based platform, cell confluence was individually measured by photo. An excel file containing individual percentage of confluence per photo and mean ± SD calculations was generated. hiPSC dishes were split every time confluence reached 70-85%.

### EB formation

EB were generated as described by Lian and Chen (2013)^20^ with minor modifications. In brief, hiPSC were resuspended in E8 medium supplemented with Polyvinyl Alcohol (PVA) and cultivated in non-adherent plates for 24 hours. Next, the medium was changed to Essential 6™ medium carefully not to remove EB in suspension. After 13 days, RNA was extracted for RT-PCR analysis.

### Karyotype

Cells were incubated with 10 μg/mL Colcemid (Sigma-Aldrich) for 1 hour and, after washing with DPBS, cells were incubated with 0.075 M KCl for 20 minutes at 37 °C. Fixation was performed by using methanol/glacial acetic acid solution (3∶1). Conventional chromosome analysis was performed on hiPSC cultures, using GTG banding at a 400-band resolution according to standard protocols with minor modifications.^21^ A total of 10 metaphase cells were analyzed. Cell images were captured using the CytoVysion system (Applied Imaging Corporation, USA).

### Integration PCR, RT-PCR and RT-qPCR

To verify if there was any episomal integration into host DNA we performed an integration PCR analysis using three set of primers (S1 Table) targeting specific sites of the plasmids DNA as described by Chou and colleagues^17^. To evaluate gene expression, RT-PCR and quantitative RT-PCR (qPCR) were performed using RNA extracted from all clones at specific passages. Detailed information about the primers can be found in supplementary material (S2 Table). Further, human embryonic stem cells (hESCs) BR1 cell line ^18^ were used as positive control of pluripotency and human skin fibroblasts (the somatic cells of origin for PC4 clones) were used as negative control of pluripotency.

### Directed differentiation into keratinocytes

iPSCs were plated with mitomycin C-inactivated 3T3 cells (donation from Prof. Monica Mathor’s laboratory) at 20% confluency. After 2 days, defined-KSFM medium (Thermo Fisher, USA) supplemented with 10 ng/mL of BMP4 (R&D Systems, USA) and 1μM retinoic acid (Sigma Aldrich, USA) was used as described by Itoh and colleagues^22^. At day 4 cells were cultivated with fresh defined-KSFM medium (Thermo Fisher, USA) for at least another 24 days. Cells were then passaged in different plates depending on the experiment and characterized by immunostaining and flow cytometry. To induce superficial-layer epithelial cells, hiPSC-derived keratinocytes (hiPSC-KCs) were subjected to a 1 μM CaCl treatment for 5-7 days as previously described by Bikle and colleagues^23^.

### Directed differentiation into cardiomyocytes

hiPSCs were differentiated using a monolayer directed differentiation method modified from previous reports^17,19^ and grown in feeder-free conditions until they reached 60–70% confluence. Cells were singularized, counted and plated (2.5×10^5^ cells/cm^2^) with E8 with 5 μM of Y-27632 (Cayman Chemical, USA). E8 medium was changed daily until cells reach 100% confluence (Day 0). Medium was then changed to RPMI supplemented with B27 (Thermo Fisher, USA) without insulin (RB-) 4 μM CHIR99021 (Merck Millipore Sigma, USA). After 24 hours, medium was changed to RB-supplemented with 10 ng/mL BMP4 (R&D Systems, USA). At day 2, medium was changed to fresh RB-supplemented with 2.5 μM KY2111 and XAV939 (both from Cayman Chemical, USA). From day 4 on, medium was changed every 48 hours to fresh RPMI supplemented with 213 μg/mL Ascorbic Acid (Sigma Aldrich, USA), 500 μg/mL DPBS, 35% BSA and 2 ug/mL Plasmocin (InvivoGen, USA). Cells were cultivated for 30 days under the described conditions before passaging as single cells to specific experiments.

### Flow cytometry and Immunofluorescence

Protein expression was analyzed by Flow Cytometry (FC) and Immunofluorescence (IF). Detailed information about the antibodies can be found in supplementary material (S3 Table). For IF, all images were generated in EVOS FL (Thermo Fisher, USA). As for FC, data was acquired using Canto BD equipment and analyzed by FlowJo Software considering 1-2% of false positive events.

### Statistical Analysis

Statistical analyses were performed using GraphPad Prism 5 (USA). All descriptive data is presented as the mean ± standard deviation (SD), groups were compared using one-way ANOVA combined with Tukey’s post-hoc test and p <0.05 was considered as statistically significant.

## RESULTS

### Single-cell passaging do not affect growth rate of hiPSC providing predictable conditions for long-term cultivation

To verify any spontaneous integrations of the vectors used for reprogramming, a set of primers specific to each vector^17,24^ was used for a PCR analysis (each clone at passages 5 and 11). Only positive control samples (vector DNA) displayed expression of the expected fragments after PCR indicating no DNA integration into hiPSC clones (S2 Figure.).

hiPSC were cultivated using E8 and E8flex media on GELTREX-coated plates, where cells grew in monolayer maintaining an undifferentiated ESC-like morphology (Figure 1B). hiPSC were passaged as single-cells by enzymatic dissociation and posterior seeding with ROCK inhibitor (Y-27632), which provided a quite controllable weekly routine (Figure 1A) based on predictable growth rates once optimal seeding density was established. To validate optimal seeding density for each one of the clones, we tested different cell seeding densities (data not shown) and tracked the growth rates by monitoring the cellular confluence with CellCountAnalyser (Figure 1C; S8 Figure). hiPSCs were dissociated every time it reached 70-85% confluency, which is the logarithmic phase of cellular growth. As a result, optimal seeding densities were then established at 40,000 or 20,000 cells per cm^2^ for ACP5 and at 43,000 or 22,000 cells per cm^2^ for PC4 clones for passages every 3 or 4 days (Mondays and Thursdays passages, respectively; Figure 1A–B).

After long term cultivation, hiPSC population doubling time (PDT) was calculated for each clone to assess growth rates. No changes were detected over time, as PDT analysis showed that single-cell passaging had no significative impact in growth rates over 50 passages for none of them. (Figure 1D). In agreement with the experiment of seeding densities, we have found mean PDT for ACP5 (28.3 ± 8.5 h) was indeed statistically different (*P<0.05) compared to PC4.3 (33.0 ± 11.4 h), PC4.5 (33.9 ± 12.6 h) or PC4.6 (32.8 ± 12.5 h), confirming different growth rates between ACP5 and PC4 clones (Figure 1E).

### Long-term single-cell passaging does not affect hiPSC pluripotency or elicit chromosomal aberration

After successive single-cell passages, cells were characterized for their pluripotency markers. To assess gene expression, we measured markers NANOG, OCT4, SOX2, LIN28 and DNMT3B by RT-qPCR using hESC lineage BR1 and human-skin fibroblasts cells as positive and negative controls of pluripotency, respectively (Figure 2A). As expected, the four hiPSC clones expressed similar amounts of all five pluripotency markers compared to BR1. Additionally, these cells also showed significantly higher levels of expression than human-skin fibroblasts (Figure 2A). Protein expression was also verified by immunostaining which confirmed that all clones expressed pluripotency markers OCT4, NANOG, and TRA-1-60 (Figure 2B; S3 Figure).

**Figure 2.**
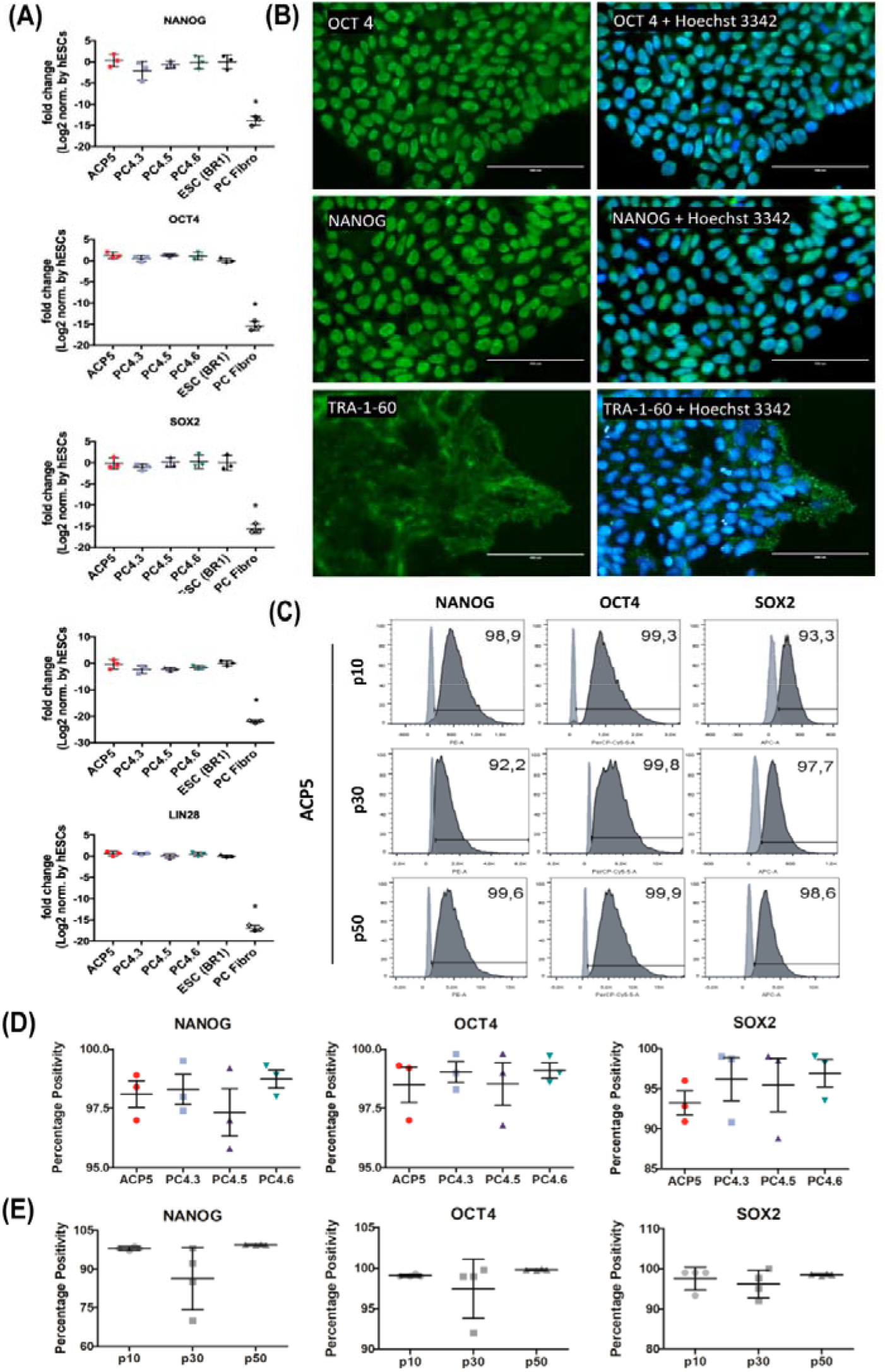
Immunocytochemistry, flow cytometry and RT-qPCR confirmed both protein and genetic expression of pluripotency markers in hiPSC clones PC4.3, PC4.5, PC4.6 and ACP5. **(A)** Expression of pluripotency markers NANOG, OCT4, SOX2, DNMT3B and LIN28 in hiPSC clones at high passages, determined by RT-qPCR. The gene expression of the hiPSC was normalized to that of BR1 (hESC). PC4 clones were obtained by reprogramming fibroblast cells (PC fibro), which were used as negative control for pluripotency. Data is presented as mean ± SD, n=3. *p< 0.05 by ANOVA with Tukey’s post hoc test. **(B)** Immunostaining of ACP5 clone at passage 20 showing expression of pluripotency markers OCT4, NANOG (both nuclear) and TRA-1-60 (membrane). Experiments were performed for all hiPSC clones **(C-E)** Cytometry data of clones at passages 10, 30 and 50 using NANOG, OCT4 and SOX2 markers. In C, light and dark grey indicate negative control (fibroblasts) and stained cells (iPSC), respectively. Statistical analysis revealed percentage positivity was not different for any marker when comparing clones in D (n = 3) or passages (n = 4) in E.

Further, NANOG, OCT4 and SOX2 were also assessed by flow cytometry using clones at passages 10, 30 and 50 to keep track of the clone’s phenotypes over time (Figure 2C; S4 Figure). Results were then analyzed for each marker in two distinctive situations, the first to compare the gene expression between different clones at the same passage, and the second to check gene expression changes over time when comparing the same clone at different passages. We observed no difference between clones (over 95% NANOG+ cells and OCT4+ cells, and over 85% SOX2+ cells, Figure 2D) neither between passages (over 70% NANOG+ cells, and over 90% OCT4+ and SOX2+ cells, Figure 2E) for any of the markers. This result suggests that not only clones are likely to display a very similar profile under same culture conditions but also that long-term cultivation had not significant impact on cells pluripotency. Chromosomal abnormalities were checked by performing a G-banding karyotype, which confirmed no aberrations at passages 10, 30 or 50 for clones ACP5, PC 4.3, PC 4.5 and PC 4.6 (Figure 3).

**Figure 3.**
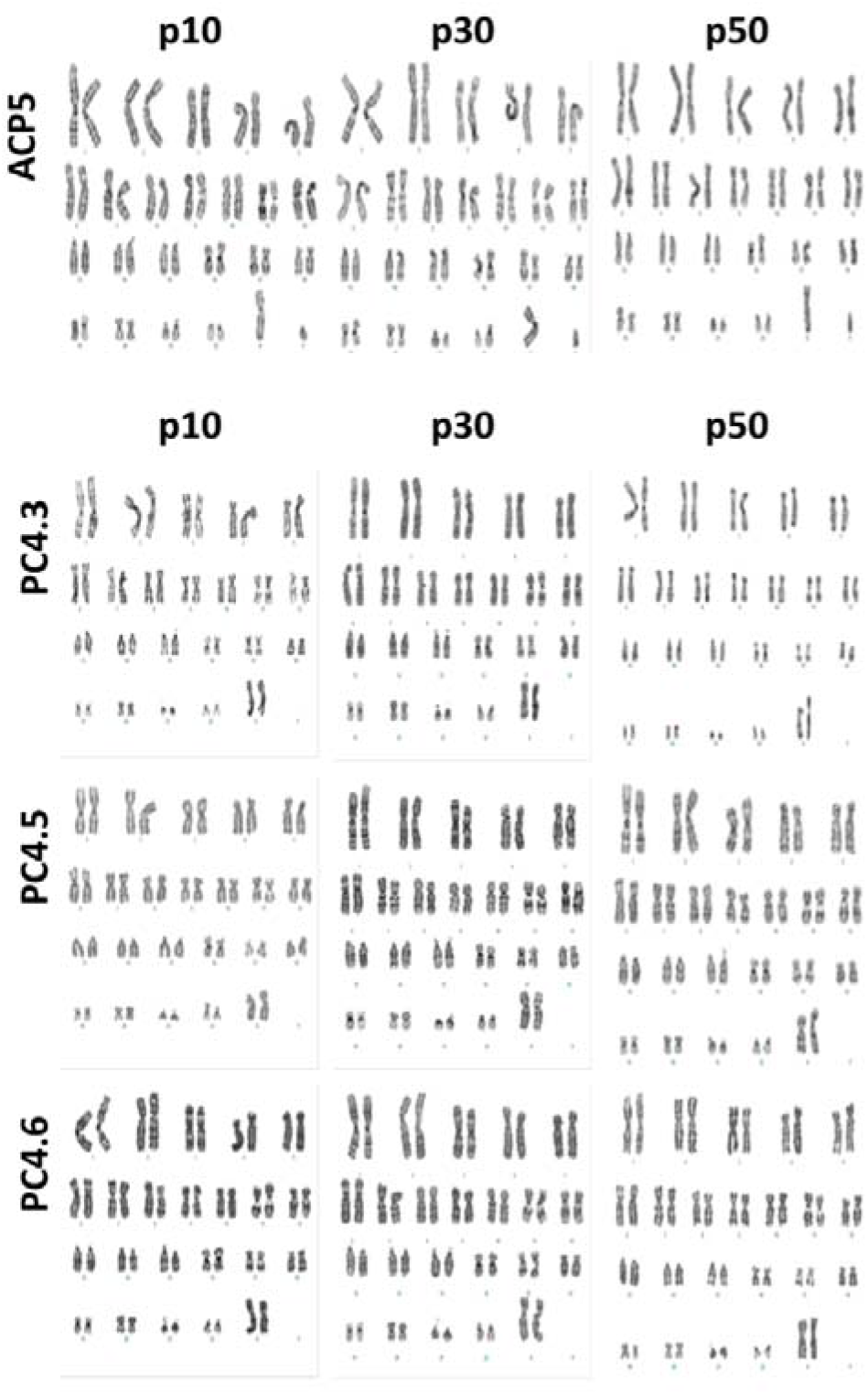
Karyotype confirmed absence of karyotypic abnormalities. G-Banding karyotype of ACP5 and PC4 clones after 10, 30 and 50 single-cell passages. No aneuploidies or translocations were detected.

### hiPSC clones subjected to single-cell passing preserve their trilineage capacity displaying high-efficiency for differentiation

The plasticity of hiPSC was tested in vitro by culturing each clone under conditions to promote embryoid body (EB) formation. As expected, after 20 days EBs revealed positive gene expression for markers such as HNF4A, MSX1 and PAX6 (endo, meso and ectoderm representants, respectively) and decreased expression of DNMT3B, a pluripotent marker (S5 Figure). In contrast, hiPSC clones at passage 20 displayed only high and expected expression of DNMT3B when in pluripotency conditions (S5 Figure).

To further evaluate the differentiation potential of long-term single-cell cultivated hiPSC, we then subjected these cells to directed differentiation protocols to generate keratinocytes and cardiomyocytes using hiPSC up to passage 70.

### Ectoderm derivative: Keratinocytes

hiPSC clones at high passages (25 to 50) were successfully differentiated into keratinocytes (hiPSC-KC). After 30 days of differentiation cells were characterized by immunostaining and flow cytometry. Flow cytometry confirmed high proportion of hiPSC-KCs positive for epithelial marker K14 (over 85%; Figure 4A) and no statistical difference of K14+ population percentage was observed between clones (Figure 4B). Immunostaining of hiPSC-KCs showed that these cells expressed not only epithelial (CD104, CD49f, K10 and K14) but also proliferative cell markers (Ki67, P63), indicating a typical profile of proliferative basal-layer epithelial cells (Figure 4C).

**Figure 4.**
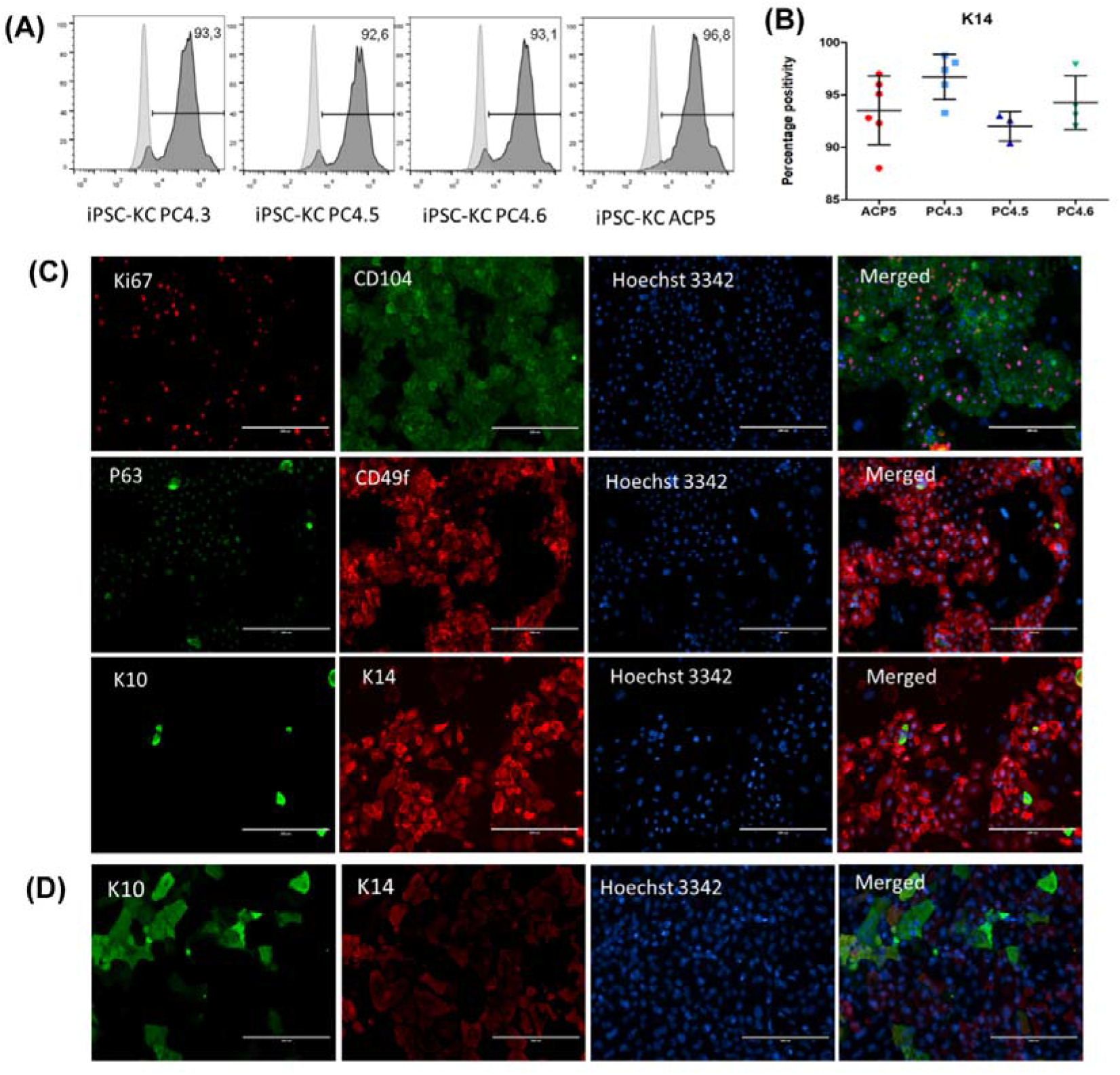
Directed differentiation into keratinocytes using hiPSC at high passages yields positive cells for both epithelial and proliferative markers. **(A-B)** Flow cytometry data showing expression of K14 marker in keratinocytes obtained from hiPSC clones PC4.3, PC4.5, PC4.6 and ACP5. Light and dark grey curves indicate negative control (hiPSC) and stained cells (hiPSC-KC), respectively. Data is presented as mean ± SD, n=3-6. **(C)** Representative immunostaining of keratinocytes obtained from PC4.3 clone (hiPSC-KC PC4.3). Ki67 and P63 are characteristic nuclear markers of proliferative cells. Surface markers CD104, CD49f and cytoskeleton markers K10, K14 are all epithelial specific. CD104, CD49f and K14 are expressed especially on proliferative basal layer cells, whereas K10 is mainly expressed on intermediate and outermost layers of epidermis. **(D)** Immunostaining of hiPSC-KC PC4.3 after calcium treatment showing higher expression of K10 and lower K14 expression. Results were confirmed by performing independent experiments for PC4.3, PC 4.5, PC4.6 and ACP5 (n = 3-6).

Additionally, we have also tested hiPSC-KC response to calcium treatment. As keratinocytes undergo the process of differentiation, cells from inner layers move up to more superficial levels and switch from producing K14 to produce K10 along with several other metabolic regulations^25^ being calcium the major regulator of keratinocyte differentiation.^23^ Thus, after treatment with calcium, hiPSC-KCs were stained using epithelial specific markers K10 and K14. Experiments were performed at least in triplicate using hiPSC-KCs derived from all clones (S6 Figure). In comparison to the hiPSC-KC without calcium treatment, we observed lower expression of K14, whereas K10 expression showed remarkable increase (Figure 4D), suggesting that long-term single-cell passaged hiPSC clones could generate keratinocytes with the ability to further differentiate into superficial-layer epithelial cells.

### Mesoderm derivative: Cardiomyocytes

We have also used hiPSC up to 70 passages for differentiation into cardiomyocytes. All hiPSC-derived cardiomyocytes (hiPSC-CMs) started to contract approximately at day 7 of the differentiation protocol. At day 30, cells were characterized by immunostaining and flow cytometry. Immunostaining showed positive expression of cardiac proteins such as NKX2-5, TNNI1, TNNI3, MYH7, TNNT2 and ACTN2 (Figure 5A). Flow cytometry showed that hiPSC-CMs derived from pluripotent cells ranging from passages 30 to 70 were generated with high-efficiency (over 80% ACTN2, TNNT2, TNNI1 and TNNI3 positive cells) and that ACP5 and PC4 lineages generated similar amounts of positive cardiac cells (Figure 5B–C). Interestingly, we have found that hiPSC-CMs exhibited simultaneous expression of TNNI1 and TNNI3, which are especially expressed in fetal and mature cardiomyocytes, respectively.

**Figure 5.**
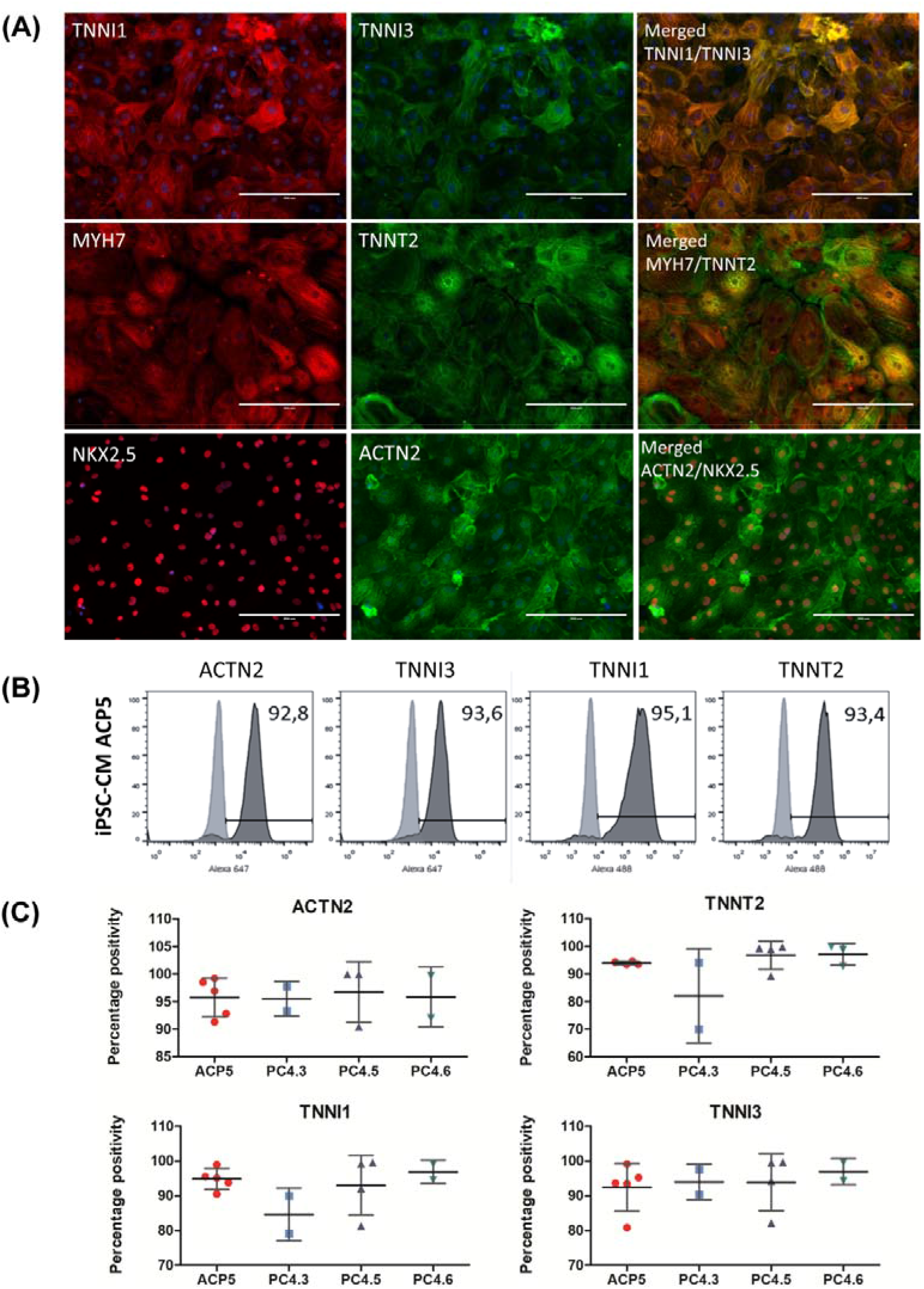
Directed differentiation into cardiomyocytes using hiPSC at high passages yields positive cells for cardiac markers. **(A)** Immunostaining of cardiomyocytes (iPSC-CM ACP5) obtained from ACP5 clone showing protein expression of cardiac specific-markers TNNI1, TNNI3, MYH7, TNNT2, NKX2.5 and ACTN2. **(B-C)** Flow cytometry data showing protein expression of cardiac-specific markers TNNI1, TNNI3, TNNT2 and ACTN2 in hiPSC-CM. Results were confirmed by performing independent experiments using APC5, PC4.3, PC4.5, PC4.6. Data is presented as mean ± SD, n=2-5.

## DISCUSSION

In the present study, we have successfully demonstrated that our cultivation method can maintain hiPSC obtained from different somatic cells and different reprogramming experiments for over 50 passages under single-cell conditions. To do so, hiPSC were characterized at different timepoints for their expression of pluripotency markers, genetic integrity, growth rates and potential to generate derivatives by both EB formation and direct differentiation.

To confirm reprogramming success, we first analyzed hiPSC for their pluripotency markers by immunocytochemistry, flow cytometry and RT-qPCR. We have found by flow cytometry that protein expression of markers OCT4, NANOG and SOX2 showed no alteration neither decrease over time for none of the hiPSC clones, corroborating permanent reprogramming as suggested by previous studies.^26,27^ Interestingly, even though reprogramming was performed using different set of plasmids and different somatic cells of origin for lineages ACP and PC4, positivity percentage for each marker showed no difference between hiPSC clones. Consistent with this, RT-qPCR also showed no significative difference among clones for gene expression of pluripotency markers *NANOG, OCT4, SOX2, LIN28* or *DNMT3B*.

Then, we analyzed the quality of the hiPSC concerning genomic alterations. As reprogramming and cultivation processes are recognized as the main causes for the alterations found in genome,^9^ genomic integrity was assessed by integration PCR and karyotyping. Because of the potential hazardous effects of reprogramming vectors,^28^ several different integration-free methods to generate iPSC were developed but, in comparison, the use of episomal plasmid vectors for reprogramming presents to its advantage an increased efficiency, as shown in previous studies.^17,24^ Indeed, by using plasmids pEB-C5, pEB-TG and Epi5, we were able to successfully generate hiPSC and PCR confirmed no vector integration into host DNA. It is important to acknowledge that integration of transgenes is not the only mechanism that is capable of altering DNA after reprogramming, however numerous reports concluded that further changes in the genome related to the vectors are mostly benign and unlikely to be threatening for research or therapy purposes,^29,30^ thus, no additional experiments to assess reprogramming-induced genomic alterations were performed.

As for culture-induced alterations, one of the main risks of prolonged cultivation is the progressive selection of genetic variants that are better adapted to *in vitro* culture environment.^14,31^ Several studies concerning ESC show that aneuploidies in chromosomes 12, 17 and X are commonly identified after long-term culture.^10,14^ As recently reviewed by Assou and colleagues,^9^ a karyotyping routine is essential for the quality assessment of PSCs as it can identify many unacceptable genomic abnormalities, which already have been reported to emerge after only 5 passages.^11^ However we have screened ACP5 and PC4 clones at passages 10, 30 and 50 and found no aberrations trough G-banding karyotype. Importantly, although we are encouraged by this result, we recognize the need for additional screening to exclude potential infra-karyotypic abnormalities such as 20q11.21 amplification^32^ or oncogenic mutations,^33^ which are also unacceptable for studies concerning hiPSC.

We have also assessed hiPSC quality concerning growth rates as PSCs proliferative log phase is required for the best results in many differentiation protocols such as cardiac ones^34^. Thus, tracking cell doubling rates during their maintenance to avoid predictable high confluence and hence cell cycle arrest is strongly desirable, and it can be easily performable by PDT calculation and cell confluence monitoring as routine of culturing. Thus, we have assessed population doubling time (PDT) and cell confluence from passage 20 to 60, a time range that corresponded to approximately 120 days of cultivation. Statistical analysis of different cultivation timepoints showed no impact in PDT for neither ACP5 nor PC4 clones. Conversely, other groups found increasing growth rates related to long-term cultivation^11,15^. This discrepancy may be explained by the fact that these findings were linked to aneuploidies found within the cells, as a strong correlation between the proportion of cell lines with abnormal karyotypes and population doubling has been previously reported in an extensive study with both ESC and iPSC^35^. Additionally, our software (CellCountAnalyser) was able to measure cell confluence quicker and more accurately than other software available^19,36–38^, giving us the ability to split cells always in log-phase of growing.

A final important finding was the maintenance of the differentiation potential after long-term cultivation, as all hiPSC clones at high passages were able to spontaneously differentiate into all three germlines through EB formation and into ecto- and mesoderm derivatives trough directed differentiation. Protein expression analysis of specific markers showed that directed differentiation consistently yielded cardiomyocytes and keratinocytes using hiPSC at high passages. It is important to state that we have also attempted directed differentiation into definitive endoderm (DE), but we were unable to do so (data not shown). In part, this can be explained by the fact that PSC differentiation potential can vary according to the cell type of origin,^39^ which can be reflected upon a diminished susceptibility for differentiation pathways such as DE. Herein, keratinocytes and cardiomyocytes, which are our main focus of our research, displayed high purity of cardiac and epithelial cells independently of the passage numbers or hiPSC lineage, as routinely reported for others.^13^

These facts bring two very important points of discussion concerning impact of iPSC management on differentiation success. First, low passages should not be used for directed differentiation as early-passage hiPSC may retain transient epigenetic memory from adult somatic cell sources impairing PSC plasticity^39^. Second, despite being a popular methodology for differentiation *in vitro*, it is known that EB formation is a process that shows very low reproducibility and often ends up with low purity of the desired differentiated cells^40,41^, whereas directed differentiation approaches tend to be more efficient and reliable to obtain specialized cells. This gives great advantage to single-cell passaging methodologies since they facilitate hiPSC directed differentiation by promoting seeding homogeneity and the formation of loosely packed clusters that leads to a more efficient cellular response to signalling molecules.^3^ Adaptation into single-cell culture was found to be a crucial step for differentiation into lung epithelia^42^ for example, and similarly, seeding density has been pointed out as one of the major optimizable factors for cardiomyocyte differentiation.^43^ In this sense, the use of a software to measure cell confluence, such as CellCountAnalyser, is a valuable tool for increasing the reproducibility and predictability of differentiation protocols.

Nonetheless, it is well recognized that single-cell passaging is usually followed by great loss of cellular viability^7^ and hence several reports suggest it promotes rapid selection of genetically abnormal clones that display increased survival rates. The increasing use of ROCK-inhibitor Y-27632^8^ to promote PSC viability after enzymatic dissociation also concern scientists because the lack of reported data on its long-term effects. Currently the studies addressing this question found no direct effect of Y-27632 on ESC genomic integrity^11^. While several studies found chromosomes 17 and 12 to be especially sensitive to adaptation into single-cell passaging^10,11,44^, many others did not detect such abnormalities upon long-term culture^45–47^. It has been previously suggested that the conflicting data concerning impact of single-cell passaging may be explained by this generical designation encompassing different passaging methods, such as EDTA, dispase, TryPLE and trypsin.^11,45^ However, as there are multiple different variables among studies, impact of single-cell passaging techniques remains subject for further investigation.

## CONCLUSION

Here we presented an easy long-term and single-cell passaging pipeline to cultivate hiPSCs that maintained their characteristics and karyotype under feeder-free conditions that allows a robust hiPSC cultivation routine by regulating cell numbers in a density- and time-dependent manner. Cells cultivated by this pipeline were able to differentiate into the three germlines trough EB formation and by high-efficient directed differentiation into keratinocytes and cardiomyocytes. Together our finding supports the long-term cultivation of hiPSCs in single-cell conditions for further use in cell-modeling and therapy.

## ACKNOWLEDGMENTS

We thank Professor Alexander Henning Ulrich for using his FACs equipment. Also, Professor Monica Mator and Lygia Veiga Pereira for the donation of 3T3 and hESC BR1 cells, respectively.

